# In depth sequencing of a serially sampled household cohort reveals the within-host dynamics of Omicron SARS-CoV-2 and rare selection of novel spike variants

**DOI:** 10.1101/2024.11.21.624722

**Authors:** Emily E. Bendall, Derek Dimcheff, Leigh Papalambros, William J. Fitzsimmons, Yuwei Zhu, Jonathan Schmitz, Natasha Halasa, James Chappell, Emily T. Martin, Jessica E. Biddle, Sarah E. Smith-Jeffcoat, Melissa A. Rolfes, Alexandra Mellis, H. Keipp Talbot, Carlos Grijalva, Adam S. Lauring

## Abstract

SARS-CoV-2 has undergone repeated and rapid evolution to circumvent host immunity. However, outside of prolonged infections in immunocompromised hosts, within-host positive selection has rarely been detected. The low diversity within-hosts and strong genetic linkage among genomic sites make accurately detecting positive selection difficult. Longitudinal sampling is a powerful method for detecting selection that has seldom been used for SARS-CoV-2. Here we combine longitudinal sampling with replicate sequencing to increase the accuracy of and lower the threshold for variant calling. We sequenced 577 specimens from 105 individuals from a household cohort primarily during the BA.1/BA.2 variant period. There was extremely low diversity and a low rate of divergence. Specimens had 0-12 intrahost single nucleotide variants (iSNV) at >0.5% frequency, and the majority of the iSNV were at frequencies <2%. Within-host dynamics were dominated by genetic drift and purifying selection. Positive selection was rare but highly concentrated in spike. Two individuals with BA.1 infections had S:371F, a lineage defining substitution for BA.2. A Wright Fisher Approximate Bayesian Computational model identified positive selection at 14 loci with 7 in spike, including S:448 and S:339. We also detected significant genetic hitchhiking between synonymous changes and nonsynonymous iSNV under selection. The detectable immune-mediated selection may be caused by the relatively narrow antibody repertoire in individuals during the early Omicron phase of the SARS-CoV-2 pandemic. As both the virus and population immunity evolve, understanding the corresponding shifts in SARS-CoV-2 within-host dynamics will be important.

## Introduction

As SARS-CoV-2 continues to circulate, population immunity from infections and vaccinations has resulted in the evolution of new variants that quickly become the dominant circulating strain^1,2^. This has contributed to decreased vaccine effectiveness, and in response, multiple reformulations of the SARS-CoV-2 vaccines^3–5^. The continual evolution of SARS-CoV-2 as a result of selection from the host adaptive immune system is likely to continue. Similar to this global antigenic drift, partial immunity from previous exposure may lead to the selection of new antigenic variants within hosts^6,7^. Because all variation originates from intrahost processes, understanding within-host dynamics is crucial to understanding the evolutionary trajectory of SARS-CoV-2.

To date, there has been limited evidence of positive selection of immune escape variants within individuals with acute, self-limited SARS-CoV-2 infections. We and others have found that SARS-CoV-2 infections exhibit low genetic diversity and few *de novo* mutations that reach significant frequencies^8–11^. Select studies have identified spike variants in sites known to confer antibody resistance^8,11^. Additionally, Farjo *et al.* found nonsynonymous intrahost single nucleotide variants (iSNVs) to be enriched in individuals who had been vaccinated or previously infected^11^. Regions of within-host positive selection in non-spike regions have also been detected when comparing intrahost diversity of synonymous and nonsynonymous variants (p_N_/p_S_)^12^. However, genetic hitchhiking (i.e., changes in a mutation’s frequency as a result of selection on a linked site on the same genome/chromosome) and genetic drift make it difficult to accurately detect positive selection with viruses from only a single timepoint^13^.

Most studies of serially sampled individuals come from prolonged infections in immunocompromised patients, where immune escape variants have repeatedly been found^14–18^. Prolonged infections release the virus from the frequent population bottlenecks characteristic of acute infections, increasing the amount of genetic variation and allowing time for selection to occur^19^. The selection pressures in immunocompromised individuals may differ from those in immunocompetent individuals with acute infections, with selection for increased cell-cell transmission and viral packaging^17^. Additionally, monoclonal antibodies commonly used to treat immunocompromised individuals may exert more targeted selection than a polyclonal response from prior exposure in immunocompetent individuals^20^.

To more thoroughly examine the role of positive selection within hosts during acute SARS-CoV-2 infections, we studied individuals from a case-ascertained household cohort, in which nasal swab specimens were collected daily for 10 days after enrollment. All specimens were sequenced in duplicate, allowing for robust variant calling at a very low frequency threshold (0.5%). With serial sampling and low frequency variant calling, we were able to define the within-host divergence of SARS-CoV-2 populations, detect genetic hitchhiking, and identify rare, but potentially significant, instances of positive selection in spike.

## Methods

### Cohort and Specimens

Households were enrolled through the CDC-sponsored Respiratory Virus Transmission Network – Sentinel (RVTN-S), a case ascertained household transmission study coordinated at Vanderbilt University Medical Center. All individuals provided written, informed consent and those included in the current study were enrolled in Nashville, TN from September 2021 to February 2022. The study was reviewed and approved by the Vanderbilt University Medical Center Institutional Review Board (see 45 C.F.R. part 46.114; 21 C.F.R. part 56.114). Index cases (i.e. the first household members with laboratory-confirmed SARS-CoV-2 infection) were identified and recruited from ambulatory clinics, emergency departments, or other settings that performed SARS-CoV-2 testing. Index cases and their households were screened and enrolled within 6 days of the earliest symptom onset date within the household. Vaccination status was determined by plausible self-report (report of a manufacturer and either a date or location) or vaccine verification through vaccination cards, state registries, and medical records. Only vaccines received more than 14 days before the date of the earliest symptom onset in the household were considered.

Nasal swabs specimens were self- or parent-collected daily from all enrolled household members during follow-up for 10 days and tested for SARS-CoV-2. Nasal swabs were tested by transcription mediated amplification using the Panther Hologic system. All available specimens were processed for sequencing as described below.

### Sequencing and Variant Calling

SARS-CoV-2 positive specimens with a cycle threshold (Ct) value ≤32 were sequenced in duplicate after the RNA extraction step. RNA was extracted using the MagMAX viral/pathogen nucleic acid purification kit (ThermoFisher) and a KingFisher Flex instrument. Sequencing libraries were prepared using the NEBNext ARTIC SARS-CoV-2 Library Prep Kit (NEB) and ARTIC V5.3.2 primer sets. After barcoding, libraries were pooled in equal volume. The pooled libraries (up to 96 specimens per pool) were size selected by gel extraction and sequenced on an Illumina NextSeq (2×300, P1 chemistry).

For the first specimen with adequate sequencing, we aligned the sequencing reads to the MN908947.3 reference using BWA-mem v0.7.15 ^21^. Primers were trimmed using iVar v1.2.1^22^. Reads from both replicates were combined and used to make a within host consensus sequence using a script from Xue et al.^23^. All specimens were aligned to their respective within-host consensus sequences. Intrahost single nucleotide variants (iSNV) were identified for each replicate separately using iVar ^22^ with the following criteria: average genome wide coverage >1000x, frequency 0.005-0.995, p-value <1×10^−5^, variant position coverage depth > 400x. We also masked ambiguous and homoplastic sites ^24^. Specific to this study, T11075C was found at low frequencies in 48 individuals and also masked. Finally, to minimize the possibility of false variants being detected, the variants had to be present in both sequencing replicates. Indels were not evaluated. Lineages were determined with Nextclade^25^ and Pango^26,27^, based on the within-host consensus sequence.

### iSNV Dynamics and Divergence rates

We calculated the divergence rate as in Xue *et al*.^23^. Briefly, we calculated the rate of evolution by summing the frequencies of within-host mutations (non-consensus allele in first specimen) and dividing by the number of available sites and the time since the infection began. We calculated the rates separately for nonsynonymous and synonymous mutations. We used 0.77 for the proportion of available sites for nonsynonymous mutations and 0.23 for synonymous. To determine the number of available sites, we multiplied the proportion of sites available by the length of the coding sequence of the MN908947.3 reference. Because symptoms typically start 2-3 days post infection and nasal swab collection occurred after symptom onset among most individuals, we added 2 days to the time since symptom onset to obtain the time elapsed between infection and sampling^28–30^. We excluded individuals who were asymptomatic from the divergence rate analysis, as we are not able to date their infection by symptom onset (e.g. 2-3 days prior as above). Because the calculated rate of divergence varied over the course of the infection, we also calculated the rate using the specimen with the highest viral load for each individual. In addition, we used linear regression to estimate the divergence rates in individuals with multiple specimens. We calculated per-site viral divergence for each specimen. For each person, a linear regression was performed with the per specimen divergences and the days post infection. A person’s divergence rate was the slope of this regression line. The rate was calculated for the whole genome and for each gene separately.

Mann-Whitney U tests were used to determine if the number iSNV per specimen and iSNV frequencies differed by mutation type, vaccination, and age group. Kruskal-Wallace tests were performed determine if the number iSNV per specimen and iSNV frequencies differed by clade and days post symptom onset. Mann-Whitney U tests were used to determine if the divergence rate differed by vaccination and age group. Kruskal-Wallace tests were performed to determine if divergence rate differed by clade, gene and days post infection. For the linear regression method, a Kruskal-Wallace test was also performed for the number of specimens available to test if the amount of information impacted the divergence rate calculations. All analyses were conducted using R version 4.3.1.

### Analysis of selection

The study period included the Delta, BA.1, and BA.2 variant periods of the SARS-CoV-2 pandemic. For each of these clades, we looked at the lineage-defining mutations in spike of the subsequent wave (i.e. BA.1, BA.2, and BA.4/BA.5). We compared the iSNV within our specimens to these lineage defining mutations.

We also used Wright Fisher Approximate Bayesian Computation (WFABC) to estimate the effective populations size (Ne) and per locus selection coefficient (s) based on allele trajectories^31^. Generation times of 8 hours and 12 hours were used^32–34^. To maximize the number of loci used in the calculation of Ne and to avoid violating the assumption that most loci are neutral, we estimated a single Ne using all loci in which the first two time points were one day apart. 10,000 bootstrap replicates were performed to obtain a posterior distribution. A fixed Ne was used for the per locus selection coefficient simulations, with the analysis repeated for the mean Ne, and +/− 1 standard deviation estimated from the previous step. A uniform prior between s of −0.5 and 0.5 was used with 100,000 simulations and an acceptance rate of 0.01. We estimated the 95% highest posterior density intervals using the boa package^35^ in R. We considered a site to be positively selected if the 95% highest posterior density did not include 0 for all three effective population sizes.

To understand how within-host selection relates to between host selection, we used the SARS-CoV-2 Nextstrain build^36^ (nextstrain/ncov, the Nextstrain team) to examine the global frequencies of iSNV that were under positive within-host selection in our study. We also compared the selection coefficients we estimated to the selection coefficients that Bloom and Neher^37^ estimated from the global phylogeny.

### Data Availability

Raw sequence reads are available at the NCBI Sequence Read Archive, Bioproject PRJNA1159790.

## Results

There were 212 SARS-CoV-2 infected individuals enrolled from September 2021 to February 2022 in this case-ascertained household cohort. Of these, we successfully sequenced 577/825 (70%) specimens from 105 individuals. Ninety nine out of 105 (94%) individuals had multiple specimens successfully sequenced (Figure 1A, Table S1). Consistent with the viruses circulating in the United States during this timeframe, the individuals in the study were infected with Delta, BA.1, and BA.2. Depth of coverage was generally high (Figure S1) and iSNV frequency was similar between replicates (Figures 1B).

**Figure 1.**
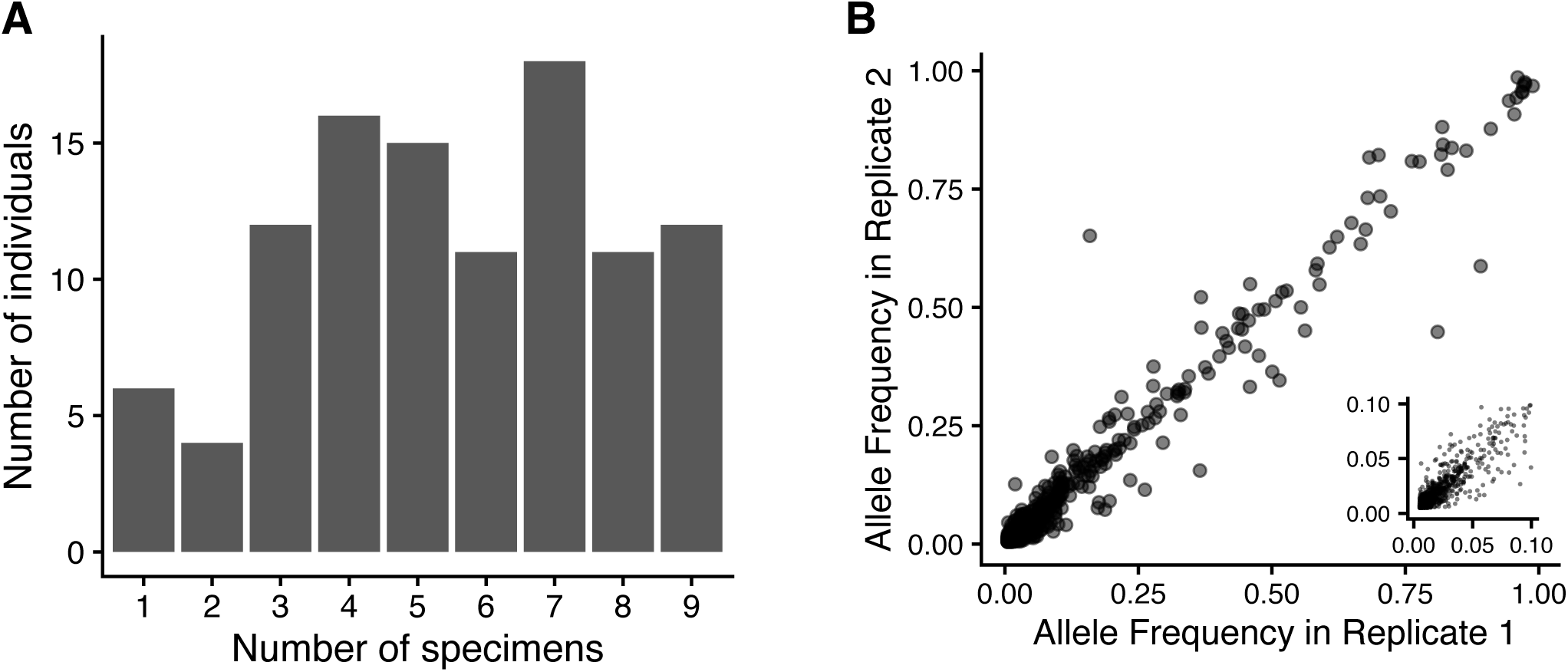
**(A)** The number of specimens per person. **(B)** intra-host single nucleotide variants (iSNV) frequency is consistent across replicates. The insert shows iSNV frequency up to 0.1

### iSNV dynamics

The allele frequencies of identified iSNV were generally very low, with the majority of iSNV present at ≤2% frequency (Figure 2A). In our cohort, the frequencies of iSNV in vaccinated individuals were higher than in unvaccinated individuals (p = 0.022, Table S2), but this difference was extremely small and unlikely to be biologically significant (Figure S2). Frequencies of iSNV also varied by the day of sampling (p = 0.002, Figure S2, Table S2) but did not differ based on host age, SARS-CoV-2 clade, or mutation type (i.e., nonsynonymous vs. synonymous; Figure S2).

**Figure 2.**
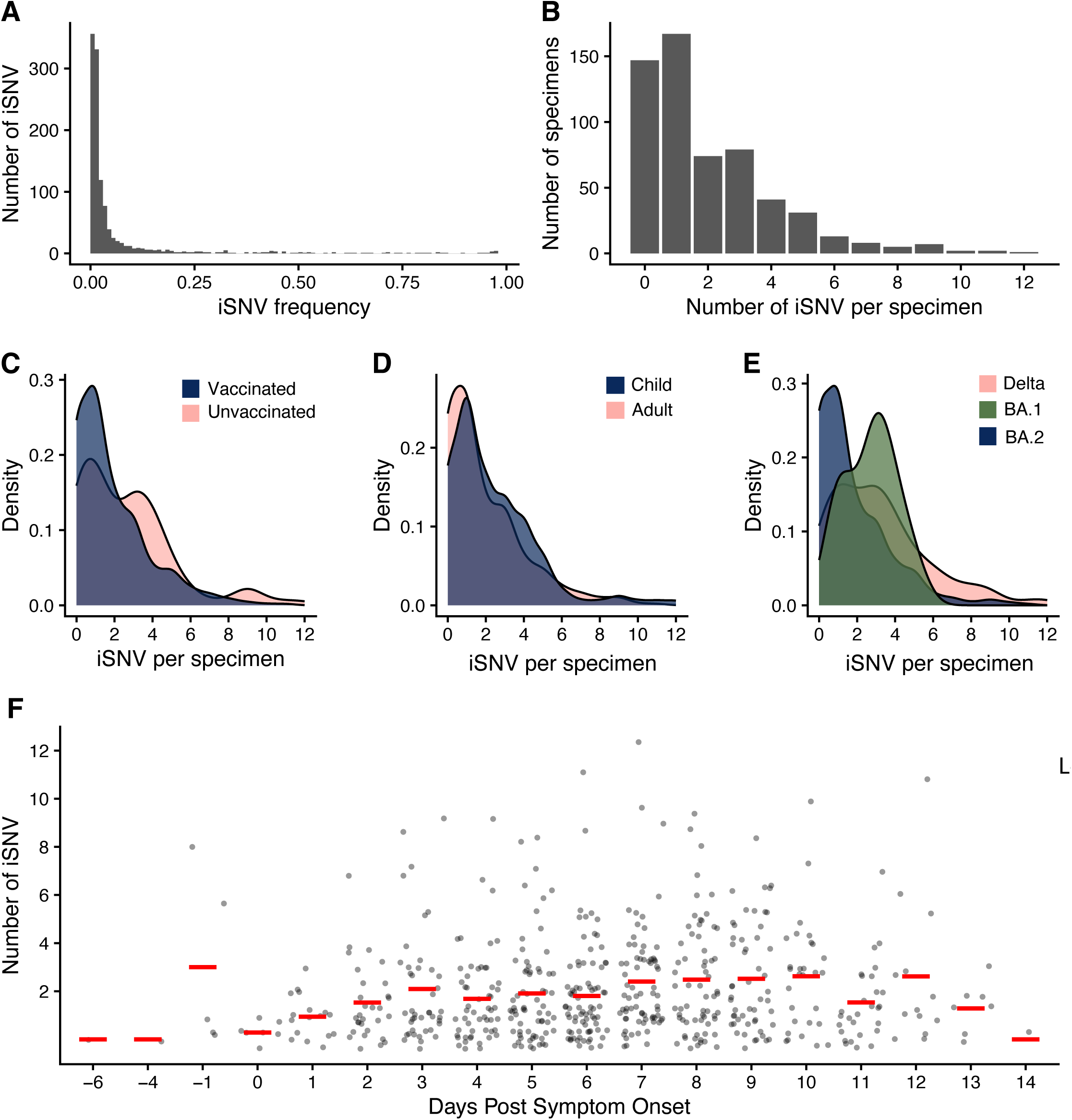
**(A)** iSNV frequency. **(B)** The number of iSNV per specimen. The number of iSNV per specimen by **(C)** vaccination status, **(D)** age with child <18 and adult 18+, **(E)** clade, **(F)** and days post symptom onset. The red lines are the mean.

All specimens had between 0-12 iSNV identified at an allele frequency ≥0.5% (Figure 2B). Unvaccinated individuals (p < 0.001) and children (p = 0.011) had greater numbers of iSNV per specimen than vaccinated individuals and adults (Figure 2C,D, Table S2). BA.1 had fewer iSNV per specimen (p < 0.001) than BA.2 (p = 0.033) or Delta (p < 0.001) infections (Figure 2E, Table S2). The number of iSNV per specimen increased as the infection progressed, and after 8-10 days post symptom onset, the number of iSNV decreased (p = 0.005, Figure 2F, Table S2). The time of sampling (days post symptom onset) did not noticeably differ by vaccination status, age, or clade (Figure S3).

### Within-host divergence rates

We estimated within-host evolutionary rates as nucleotide divergence per site per day on a per-specimen basis and by linear regression in individuals for whom we had multiple sequenced specimens. The genome-wide mean divergence rate was 5.03 x 10^-7^ nucleotide substitutions/site/day for nonsynonymous mutations and 1.08 x 10^-6^ for synonymous mutations. Although not statistically significant, the estimated divergence rate varied according to the day of sampling when using the point method (Figure 3). The divergence rate increased from the onset of the infection until approximately day 5 for nonsynonymous sites and day 8 for synonymous sites and then decreased. For the rest of the comparisons using the point method, the divergence rate from the specimen with the highest viral load was used. Children had higher rates for nonsynonymous mutations, but not synonymous mutations (p= 0.019, Figure 3C, Table S3), while rates for synonymous mutations were not associated with age. The divergence rate did not differ by vaccination status or clade (Figure 3, Table S3). There were significant differences in divergence rate based on gene (p<0.001); notably, spike had a higher divergence rate compared to ORF1a for nonsynonymous mutations, but did not differ from any of the other genes (Figure 3E,F, Table S4).

**Figure 3.**
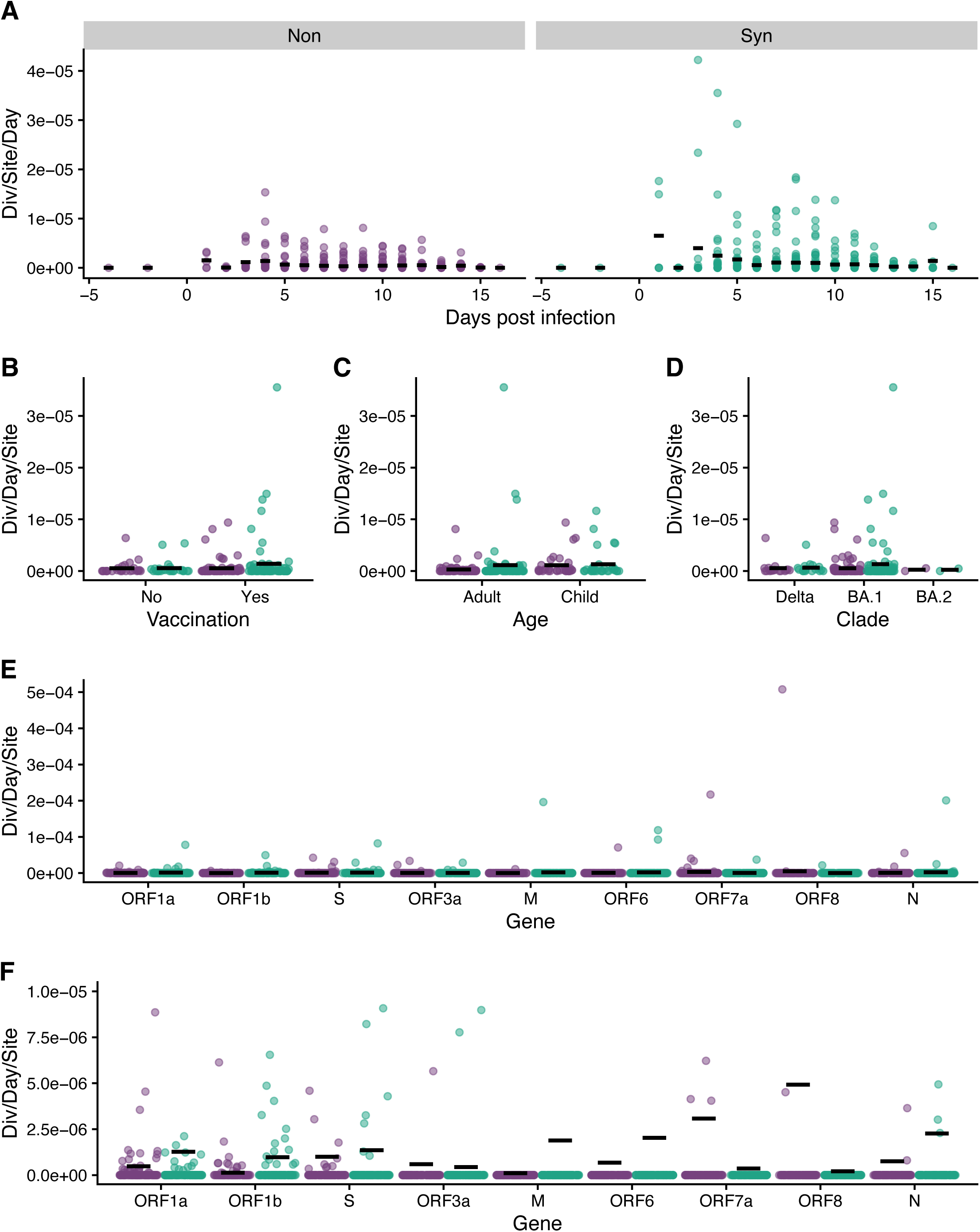
**(A)** Divergence rate (divergence/site/day) for all specimens by days post infection. Divergence rate (divergence/site/day) using the specimen with the highest viral titer by **(B)** vaccination status, **(C)** age, **(D)** clade, and **(E)** gene. **(F)** is a zoomed in version of **(E)**, note y-axis. Black lines are the mean divergence rate. Green is synonymous, and purple is nonsynonymous.

Results obtained by linear regression were slightly different. The divergence rate did not differ by vaccination, age, clade, or gene (Figure S4, Table S3). For synonymous mutations, individuals with two specimens had a lower rate than individuals with more than two specimens (p = 0.046, Figure S4, Table S3). In many cases in which there were only two specimens for an individual, these were collected after the peak of infection giving the regression a negative slope.

### Analysis of selection

We analyzed selection by first looking for iSNV that anticipated mutations that defined subsequent variants. Two individuals with BA.1 had an iSNV that causes S:371F, a BA.2 lineage defining mutation (Table 1). These iSNV were at low frequencies, with a maximum observed frequency of 0.8% and 1.8%. There were 3 additional iSNV in the codon for a lineage defining mutation but resulted in a different amino acid substitution. This included a third iSNV at position 371.

**Table 1.** intra-host single nucleotide variants (iSNV) that are in the position as lineage defining mutations in spike for the subsequent variant of concern wave.

| Clade | Next Wave | Lineage Defining Mut. | Observed Mutation | Individual | Max Obs. Frequency |
| --- | --- | --- | --- | --- | --- |
| Delta | BA.1 | N440K | N440Y | 102101 | 0.049 |
| Delta | BA.1 | G446S | G446V | 101201 | 0.018 |
| Delta | BA.1 | T547K | T547I | 101703 | 0.008 |
| BA.1 | BA.2 | S371F | L371I | 102601 | 0.008 |
| BA.1 | BA.2 | S371F | L371F | 105701 | 0.018 |
| BA.1 | BA.2 | S371F | L371F | 107303 | 0.008 |

Using a WFABC model, we estimated a within-host effective population size of 78. Fourteen iSNV from 11 individuals were under positive selection: 7 in spike, 6 in other coding regions and 1 in a non-coding region (Figure 4A, B, Table 2). The results were the same for 8hr and 12hr generation times. Of the iSNV found in coding regions, 10 were nonsynonymous, including 6 of the iSNV in spike. Two of the selected synonymous iSNV were in individuals that had nonsynonymous iSNV under positive selection, suggestive of linkage as the allele trajectories of the two iSNV were closely matched (Figure 4C, D).

**Figure 4.**
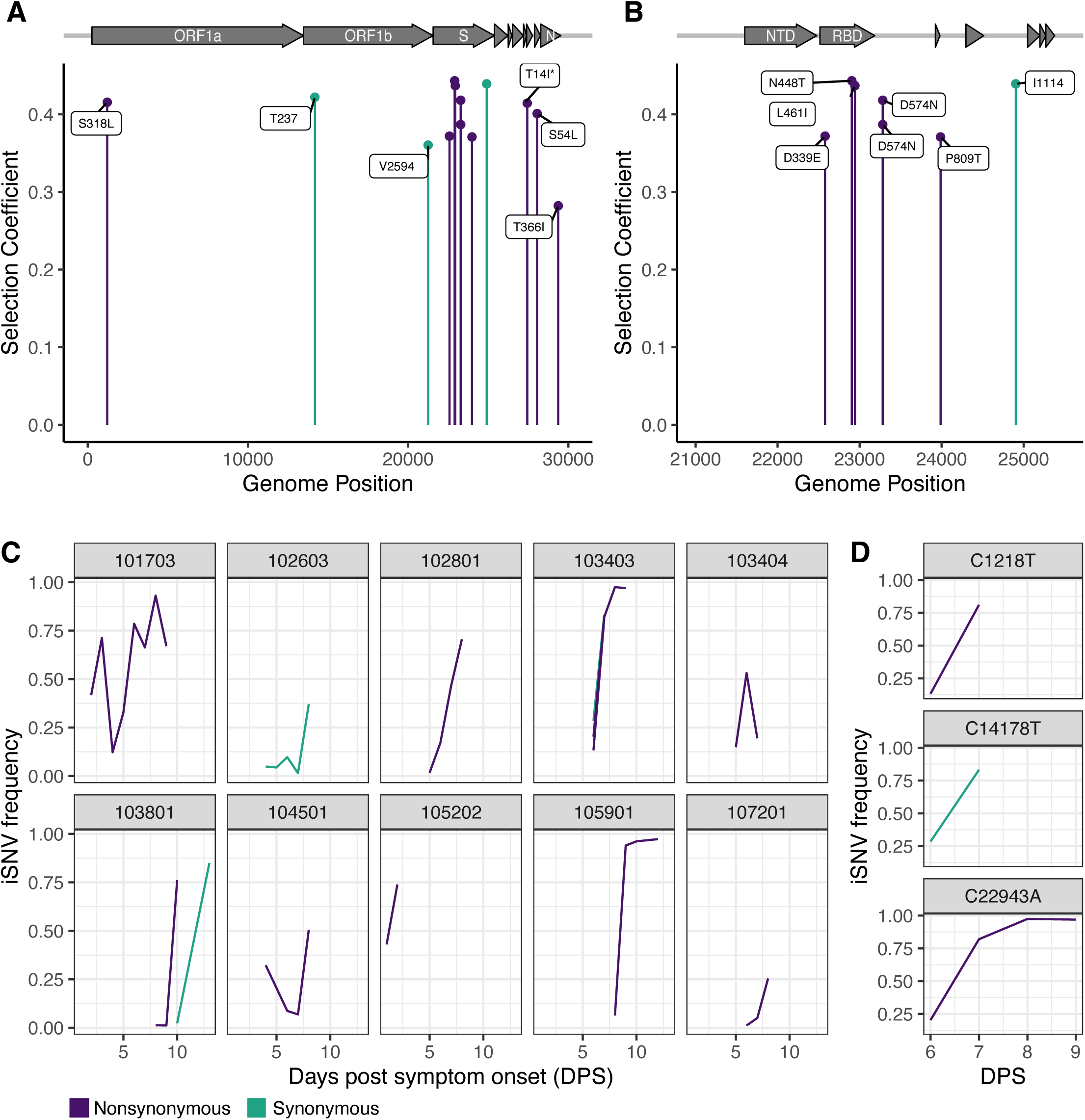
**(A)** WFABC selection coefficients for iSNV under positive selection for the whole genome and **(B)** for spike. **(C)** The allele trajectories of the iSNV with positive selection coefficients by individual. **(D)** The allele trajectory for iSNV in individual 103403, denoted with an asterisk in **(C)**. Green is synonymous, and purple is nonsynonymous. WFABC = Wright Fisher Approximate Bayesian Computation; iSNV = intra-host single nucleotide variants

**Table 2.** iSNV with positive selection coefficient. Selection coefficient values are from the 8hr generation time results.

| Gene | iSNV | AA Mutation | Mutation Type | Selection Coefficient | Individual | Age | Clade | Vaccinated |
| --- | --- | --- | --- | --- | --- | --- | --- | --- |
| ORF1a | C1218T | S318L | Non | 0.41550572 | 103403 | 42 | BA.1 | Yes |
| ORF1b | C14178T | 237T | Syn | 0.42198689 | 103403 | 42 | BA.1 | Yes |
| ORF1b | G21249T | 2594V | Syn | 0.36034801 | 102603 | 46 | BA.1 | Yes |
| S | T22579A | D339E | Non | 0.37183336 | 105202 | 54 | BA.1 | Yes |
| S | A22905C | L461I | Non | 0.43687486 | 103403 | 42 | BA.1 | Yes |
| S | C22943A | N448T | Non | 0.44314941 | 105901 | 49 | BA.1 | Yes |
| S | G23282A | D574N | Non | 0.38677793 | 103404 | 36 | BA.1 | Yes |
| S | G23282A | D574N | Non | 0.41807759 | 102801 | 39 | BA.1 | Yes |
| S | C23987A | P809T | Non | 0.37079572 | 107201 | 50 | BA.1 | Yes |
| S | C24904T | 1114I | Syn | 0.43920636 | 103801 | 47 | BA.1 | No |
| ORF7a | C27434T | T14I | Non | 0.41432335 | 103801 | 47 | BA.1 | No |
| ORF8 | C28054T | S54L | Non | 0.40091975 | 101703 | 10 | Delta | No |
| N | C29370T | T366I | Non | 0.28217821 | 104501 | 10 | BA.1 | Yes |
| NA | G29742A | NA | NA | 0.35436237 | 106103 | 36 | BA.1 | Yes |

Footnote: iSNV = intra-host single nucleotide variants

Three of the selected spike amino acid substitutions were in the RBD (Receptor Binding Domain). Outside of the RBD, two individuals shared the positively selected substitution, S:D574N. A third individual had S:D574N in 4 specimens, yet without a positive selection coefficient. None of the iSNV in future lineage defining codons had a positive selection coefficient. However, one individual had both an iSNV in a lineage defining codon (S:547) and an iSNV with a positive selection coefficient in the viral replicase (ORF8:S54L). All of the nonsynonymous spike iSNV were in vaccinated individuals.

We used the SARS-CoV-2 nextstrain build to determine whether any of the iSNV with positive selection coefficients were also identified as increasing in global frequency^36^. None of the iSNV or resulting amino acid changes reached more than 5% globally (data not shown). The selection coefficients we estimated were only weakly related to the between host selection coefficients estimated by Bloom and Neher^37^ (Figure S5).

## Discussion

In this intensive evaluation of serially sampled individuals in a longitudinal household transmission study, we found that within-host SARS-CoV-2 populations are dominated by purifying selection and genetic drift. This results in low levels of diversity and low rates of divergence, consistent with previous studies^8–10,38,39^. There were differences in divergence rate based on age and in the frequency of iSNV based on vaccination, but these are unlikely to be biologically significant. Multiple factors influenced the number of iSNV per specimen, notably day of sampling. Positive selection was rare, but when present, it tended to be enriched in spike and the RBD.

The low level of diversity is similar to what we and others have reported for SARS-CoV-2^8–10,38,39^. Some study-specific differences in diversity are noteworthy. For example, Farjo *et al.* (with specimens from 40 individuals) observed higher numbers of iSNV in vaccinated individuals, while we found higher numbers of iSNV in unvaccinated individuals^11^. However, their quality metrics differed between vaccinated and unvaccinated individuals, and their sample size was smaller than the present study. Additionally Gu *et al.* found that the number of iSNV per specimen was higher in VOC compared to non-VOC clades, but did not find any differences between VOC clades^12^. In contrast, we found Delta and BA.2 had more iSNV than BA.1. Our frequency threshold for variant calling was lower, and potentially more sensitive to differences in iSNV number. Variation between cohorts likely contributes to differences between studies, but different study designs and methods also account for dissimilarities.

SARS-CoV-2 has comparable within-host dynamics to influenza A virus. The distribution of allele frequencies is very similar in influenza A and SARS-CoV-2, with most iSNV found at very low frequencies^23,40,41^. However, compared to studies of influenza with the same iSNV threshold, SARS-CoV-2 had fewer iSNV per specimen despite the genome being twice the size. SARS-CoV-2 also had lower divergence rates of 10^-6^ div/site/day for synonymous sites and 10^7^ for nonsynonymous sites, compared to 10^-5^ and 10^-6^ for influenza A in synonymous and nonsynonymous sites respectively^23,40^. The lower within-host diversity of SARS-CoV-2 is largely attributable to the difference in mutation rates. With its proofreading capabilities, SARS-CoV-2 has a mutation rate of 9 x 10^-7^ mutations per nucleotide per replication cycle^42^ compared to 2 x 10^-6^ in influenza A (using analogous assays)^43^. The strength of genetic drift may also contribute to the observed differences. While both influenza A virus (Ne ∼150-300)^40,41^ and SARS-CoV-2 have small effective population sizes, the smaller effective population size in SARS-CoV-2 will result in stronger genetic drift. More of the variation will be lost from the population or not repeatedly sampled due to changes in population structure. These within-host dynamics are largely consistent with the neutral theory of evolution^44^. Strongly deleterious mutations are removed quickly from the population and the remaining variation is largely neutral.

Despite overall similar patterns of within-host dynamics between SARS-CoV-2 and influenza A virus, there are differences in the nature of selected sites. In influenza A virus, we have not found an overrepresentation of selected sites in hemagglutinin (HA), including antigenic sites, or in neuraminidase (NA)^40^. In contrast, in SARS-CoV-2 we found a greater number of positively selected sites in spike (7/13) and in the RBD (3) than expected by chance. This is consistent with selection for immune escape. Within the RBD, S:D339E was under positive selection. Although this exact amino acid substitution has not previously been known to be under selection, S:339 is the most variable amino acid in spike^45^. Additionally, G339D is a lineage defining mutation in BA.1, BA.2, BA.4, and BA.5^46^, and D339H is a lineage defining mutation for BA.2.75, XBB, and BA.2.86^47,48^. Both of these amino acid substitutions have been shown to escape neutralizing antibodies^49,50^.

In the RBD, S:448 is an epitope targeted by multiple monoclonal antibodies, including bebtelovimab, imdevimab, and cilgavimab^46^. These monoclonal antibodies have high similarity to germline encoded antibodies^51–53^, making S:448 an epitope that is likely to be commonly targeted across individuals. Outside of the RBD, two individuals in different households had D574N under positive selection. This substitution has been observed in a long-term infection of an immunocompromised patient^54^ and also detected in a small proportion of BA.5 lineages. This infrequent but detectable positive selection may be due to the timing of these infections relative to viral emergence. This study enrolled individuals within approximately the first 18-24 months of the pandemic. At this time, only the Wuhan strain spike was used for vaccination, leading to a relatively narrow antibody repertoire. A narrow antibody repertoire may cause uniform selection pressure, with one or a few mutations being sufficient for SARS-CoV-2 to be resistant to a majority of the host antibodies, similar to treatment with monoclonal antibodies^51,55^. In our study, six of the selected sites in spike, all of the nonsynonymous sites, and all of the selected sites in the RBD occurred in vaccinated individuals. Over time as the number of exposures and lineages individuals are exposed to increases, their antibody repertoires also increase^56,57^. As the antibody repertoire diversifies, individual mutations may make SARS-CoV-2 resistant to only a small proportion of antibodies, leading to weaker selection^57^. Earlier in the pandemic there may have been low levels of selection due to lack of even partial immunity, coinciding with a period of global evolutionary stasis^42^.

Despite finding immunologically relevant iSNV, our results had low predictive power for trends in SARS-CoV-2 evolution globally. None of the iSNV under positive selection or the corresponding amino acid substitutions reached >5% frequency globally at any time. Two individuals with BA.1 infections had a lineage defining mutation, S:371F, for BA.2. However, the mutation remained at very low frequencies within these two individuals. In the first individual, the selection coefficient was not statistically significantly different than 0 (s= −0.07), and a selection coefficient was unable to be calculated for the second individual due to the number of specimens. With low effective population sizes and stochastic dynamics, our estimates of positive selection are conservative. However, combining within-host variant data with other sources (e.g. deep mutational scanning or inferred between-host selection coefficients) may be fruitful for understanding the evolutionary trajectory of SARS-CoV-2.

A major strength of this study is daily sampling, with up to 9 successfully sequenced specimens per individual, allowing us to examine allele trajectories. Summary statistics meant to detect selection can be misleading due to genetic linkage and hitchhiking^13^. These effects are especially prominent in cases where there are strong bottlenecks and low levels of recombination. With serial sampling, we were able to calculate selection coefficients and detect hitchhiking of synonymous mutations with a physically linked nonsynonymous mutation. To illustrate, in one individual, there were three iSNV with nearly identical allele trajectories: 2 nonsynonymous and 1 synonymous. Most likely, the nonsynonymous iSNV in ORF1a and the synonymous iSNV in ORF1b were swept along with the nonsynonymous iSNV in spike (L461I).

Our study has several limitations. First, our results may not generalize to other phases of the SARS-CoV-2 pandemic. The study took place over 6 months in the second year of the pandemic after the availability of a single vaccine formulation. Results may differ as vaccine and exposure history become more variable across the population and as SARS-CoV-2 has had a longer evolutionary history with human hosts. Indeed, we speculate that SARS-CoV-2 evolution during acute infections could become more similar to the dynamics of influenza A virus within hosts^40,41^. Second, there is always the possibility of inaccurate variant calls. However, this possibility was mitigated by sequencing all specimens in replicate and sequencing multiple specimens per person reduces this possibility. Third, SARS-CoV-2 has significant compartmentalization^58,59^, and we are only sampling one location in the body; but when compared, nasal and saliva specimens have similar within-host dynamics dominated by stochastic processes^11^.

Across studies, acute respiratory viruses have similar within-host dynamics: tight bottlenecks, low genetic diversity, and populations dominated by purifying selection and genetic drift^8–12,19,38–41,60,61^. Overall, our findings are consistent with this pattern. However, nuanced differences exist between viruses, cohorts, and demographic features. In our cohort, within-host positive selection was rare, but appeared to frequently be immune mediated when present. As viruses adapt to human hosts and the population develops immunity, it will be important to follow the shifting impacts on within-host dynamics and selective pressure.

## Acknowledgements

We thank all participants in the study for their time and effort. Primary funding for the RVTN study was provided by the US Centers for Disease Control and Prevention (CDC 75D30121C11656). CGG was partially supported by NIH K24A I148459. Scientists from the US CDC participated in all aspects of this study, including its design, analysis, interpretation of data, writing the report, and the decision to submit the article for publication. Sequencing and associated analysis was supported by a NIH R01 AI148371 (to ASL and ETM), and the Penn Center for Excellence in Influenza Research and Response, Penn-CEIRR, NIH 75N93021C00015 (to ASL and ETM) and the Michigan Infectious Disease Genomics Center, NIH (to ASL).

## Disclaimer

The findings and conclusions in this report are those of the authors and do not necessarily represent the official position of the Centers for Disease Control and Prevention (CDC).

## Author Contributions

Conceptualization: Lauring, Grijalva, Talbot

Data Curation: All authors

Formal Analysis: Bendall, Zhu, Lauring

Funding Acquisition: Lauring, Martin, Grijalva, Talbot

Investigation: All authors

Methodology: All authors

Manuscript writing: Bendall, Lauring

Manuscript editing: All authors

Project Administration: Lauring, Grijalva, Talbot

Resources: All authors

---

**Figure S1.**
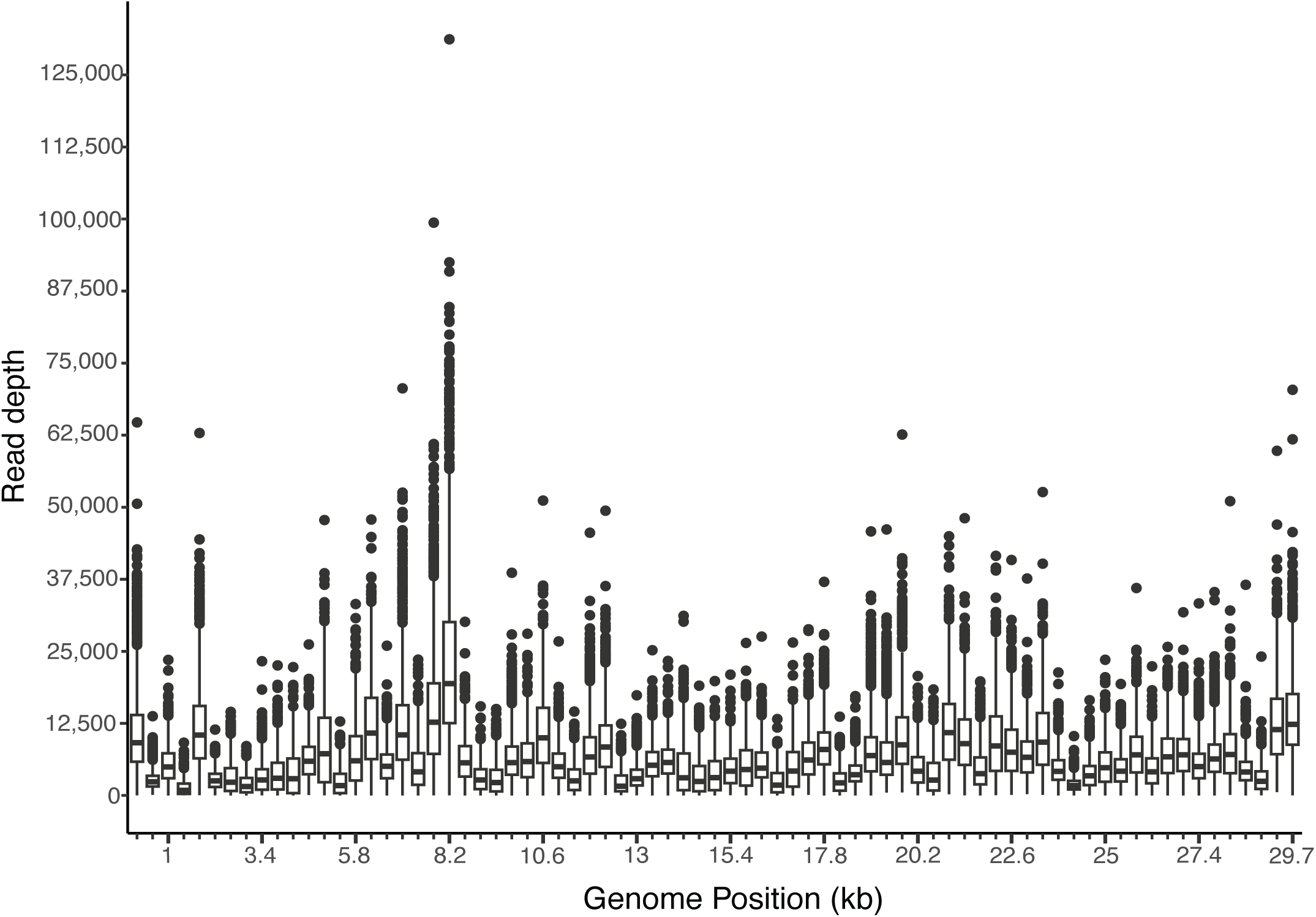
Boxplots of coverage across the genome in non-overlapping windows of 400 bp for specimens with high quality sequencing. The box shows the first quartile, median, and third quartile. The whiskers are 1.5x interquartile range, and the dots are the outliers.

**Figure S2.**
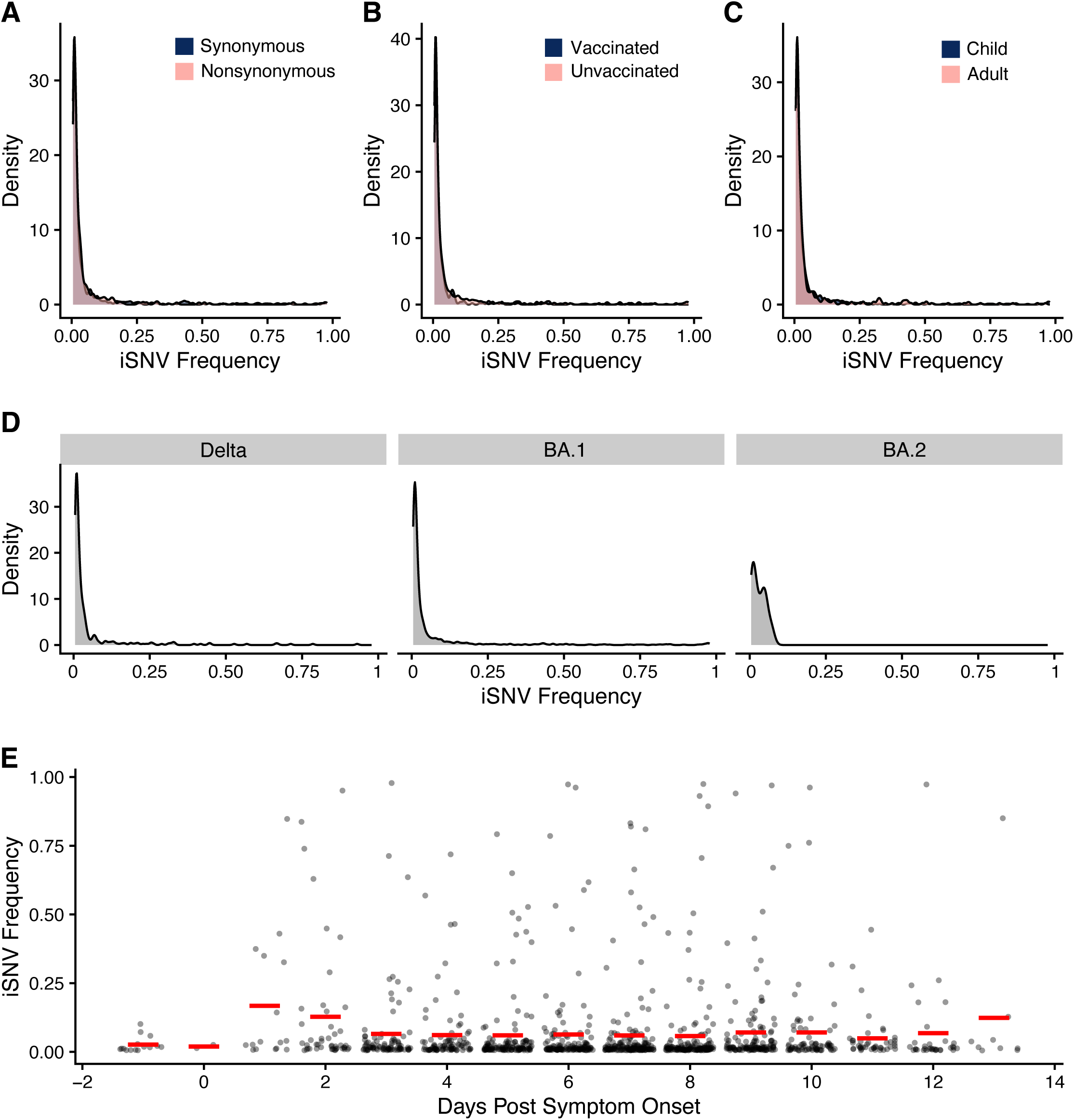
iSNV frequency by **(A)** mutation type, **(B)** vaccination status, **(C)** age with child <18 and adult 18+, **(D)** clade, and **(E)** days post symptom onset. The red lines are the mean. iSNV = intra-host single nucleotide variants.

**Figure S3.**
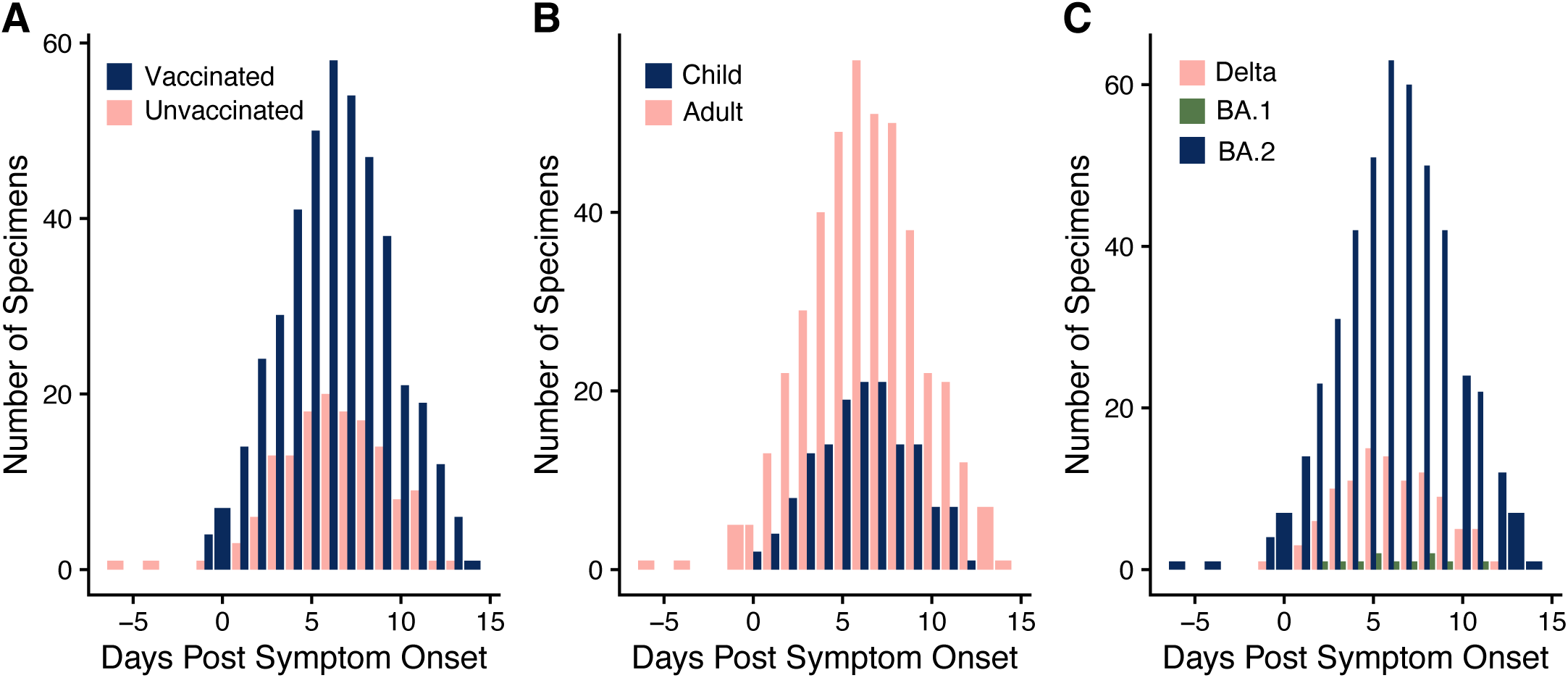
Number of specimens collected per day post symptom onset by **(A)** vaccination status, **(B)** age with child <18 and adult 18+, and **(C)** clade.

**Figure S4.**
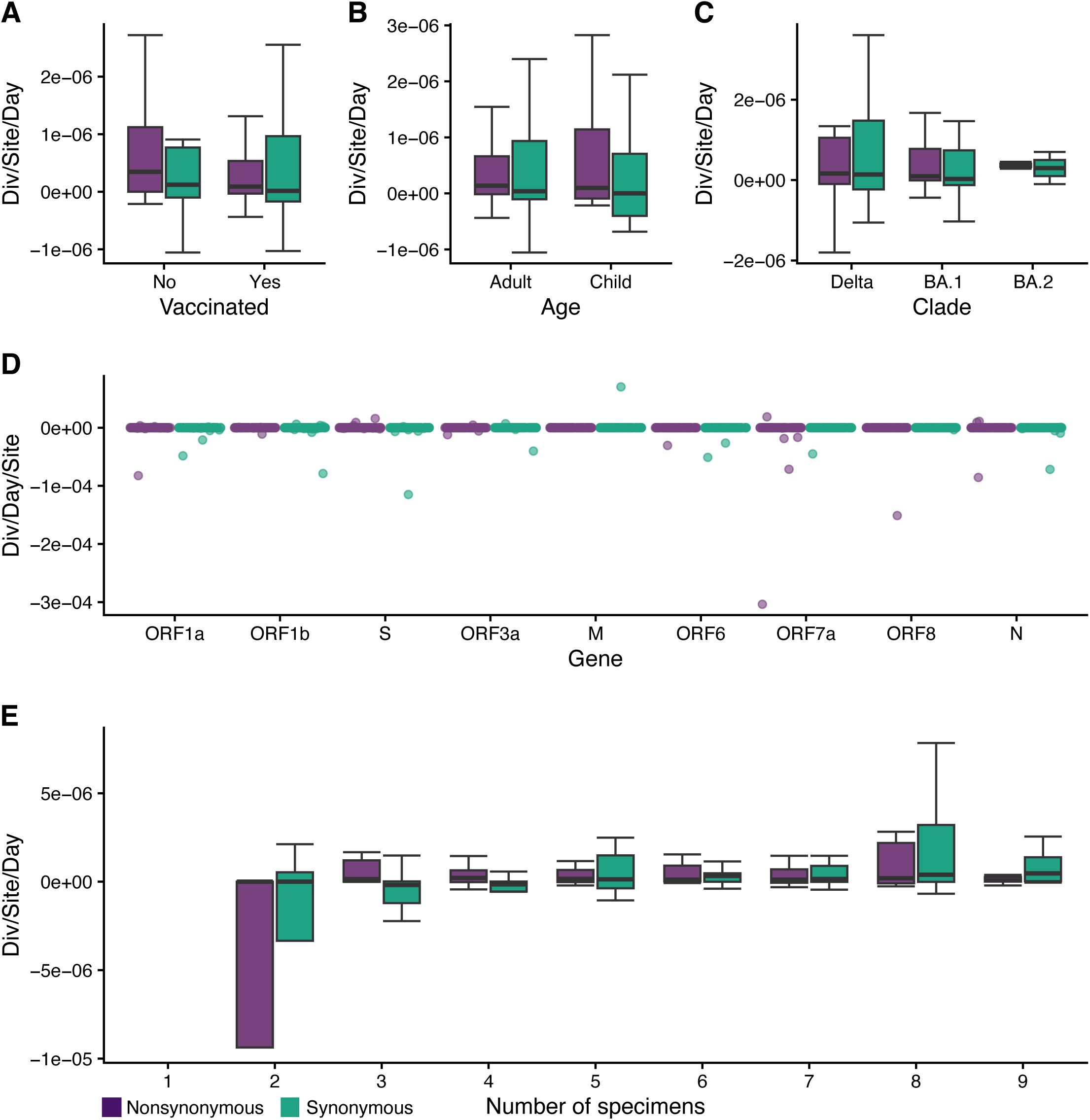
Divergence rate (divergence/site/day) using linear regressions by **(A)** vaccination status, **(B)** age with child <18 and adult 18+, **(C)** clade, **(D)** gene, **(E)** and number of specimens used in the calculation (green synonymous, purple nonsynonymous).

**Figure S5.**
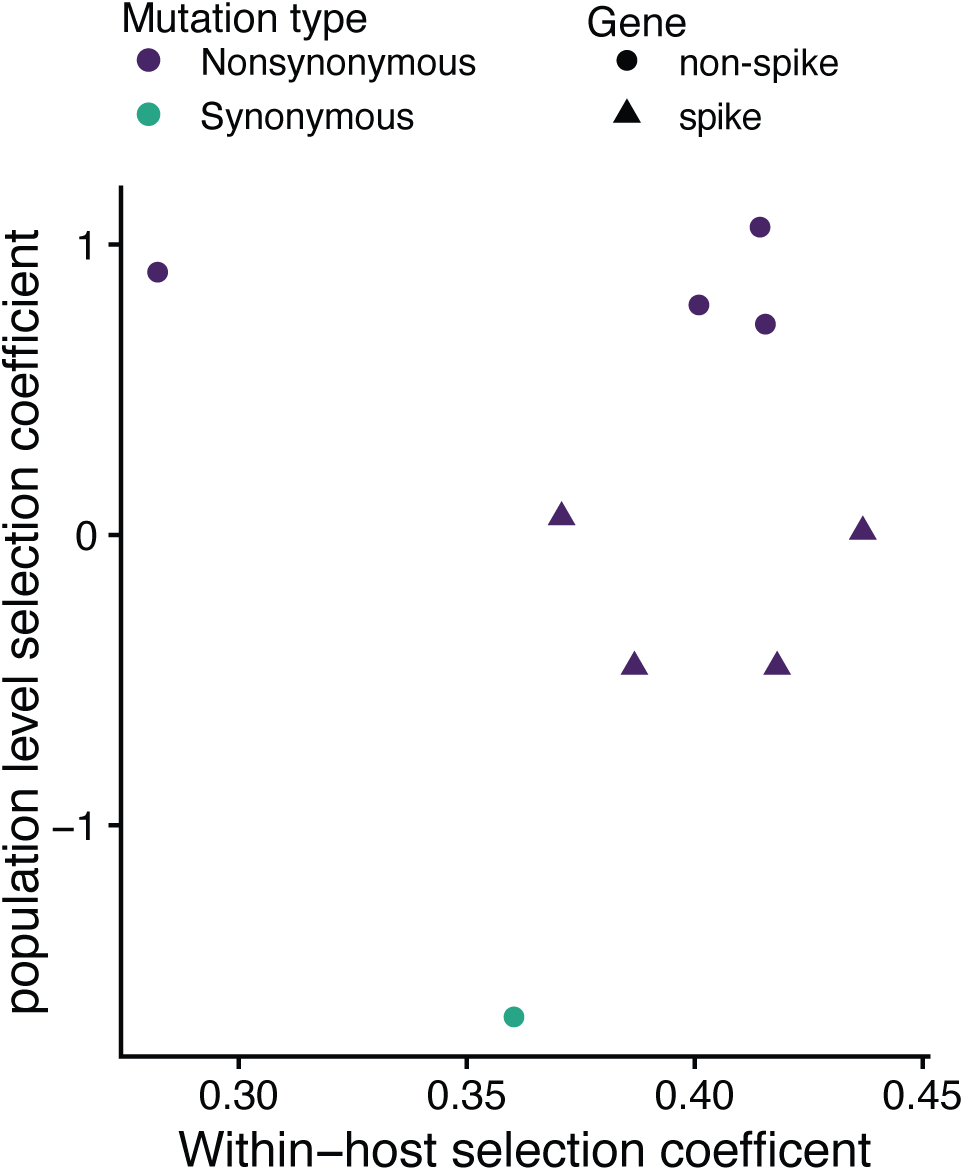
Comparison of the within-host selection coefficient and the population level selection coefficient for Bloom & Neher 2023. Green is synonymous and purple is nonsynonymous. Triangles are mutations in spike z circles are in non-spike genes.

**Table S1.** Demographic information and infection details for individuals in this study (n=105)

| Characteristic |  | n |
| --- | --- | --- |
| **Age** | Child (<18 years) | 32 |
|  | Adult (≥18 years) | 73 |
| **Clade** | Delta | 17 |
|  | BA.1 | 86 |
|  | BA.2 | 2 |
| **Vaccination** | Yes | 76 |
|  | No | 23 |
|  | Missing | 6 |
| **Sex** | Female | 57 |
|  | Male | 47 |
|  | Missing | 1 |
| **Symptomatic** | Yes | 103 |
|  | No | 1 |
|  | Missing | 1 |
| **Multiple specimens sequenced** | Yes | 99 |
|  | No | 6 |

**Table S2.** Comparisons of the number of iSNV per specimen and of iSNV frequency. For statistically significant differences the p values are bolded. *x*^2^ test statistics are from Kruskal-Wallis rank sum tests and W test statistics are from Mann-Whitney U tests.

| **Number of iSNV per sample** | Test statistic ($\chi^2$ or W) | df | p value |
| --- | --- | --- | --- |
| Age | 28066 | 1 | **0.011** |
| Vaccination | 37242 | 1 | **< 0.001** |
| Days Post Symptom Onset | 35.768 | 17 | **0.005** |
| Clade | 38.751 | 2 | **< 0.001** |
| ***Clade - Post Hoc*** | Test statistic (Z) | p.unadj | p.adj |
| Delta vs BA.1 | 5.890058 | **< 0.001** | **< 0.001** |
| Delta vs BA.2 | -0.278536 | 0.781 | 0.781 |
| BA.1 vs BA.2 | -2.392985 | **0.017** | **0.033** |
| **iSNV Frequency** | Test statistic ($\chi^2$ or W) | df | p value |
| Age | 144856 | 1 | 0.792 |
| Vaccination | 135509 | 1 | **0.022** |
| Clade | 2.4914 | 2 | 0.288 |
| Days Post Symptom Onset | 34.069 | 17 | **0.002** |

Footnote: iSNV = intra-host single nucleotide variants

**Table S3.** Comparisons of divergence rates. Statistically significant differences are bolded. *x*^2^ test statistics are from Kruskal-Wallis rank sum tests and W test statistics are from Mann-Whitney U tests.

|  | **Nonsynonymous** |  |  | **Synonymous** |  |  |
| --- | --- | --- | --- | --- | --- | --- |
| | Test statistic<br>($\chi^2$ or W) | df | p value | Test statistic<br>($\chi^2$ or W) | df | p value |
| ***Point Estimate*** |  |  |  |  |  |  |
| Gene | 76.888 | 8 | **<0.001** | 55.212 | 8 | **<0.001** |
| Age | 821.5 | 1 | **0.019** | 957.5 | 1 | 0.165 |
| Days Post Symptom Onset | 21.724 | 17 | 0.196 | 15.159 | 17 | 0.584 |
| Vaccination | 1040.5 | 1 | 0.835 | 1034 | 1 | 0.869 |
| Clade | 1.6218 | 2 | 0.444 | 3.095 | 2 | 0.213 |
| ***Logistic Regression*** |  |  |  |  |  |  |
| Gene | 7.6549 | 8 | 0.468 | 12.619 | 8 | 0.126 |
| Age | 970 | 1 | 0.930 | 1056 | 1 | 0.440 |
| Vaccination | 1057.5 | 1 | 0.239 | 980 | 1 | 0.585 |
| Clade | 0.74911 | 2 | 0.688 | 0.044783 | 2 | 0.978 |
| Number of Specimens | 3.8194 | 7 | 0.800 | 14.318 | 7 | **0.046** |

**Table S4.** Post hoc (Dunn) tests for divergence rate between genes using the point.

| Nonsynonymous |  |  |  |  | Synonymous |  |  |
| --- | --- | --- | --- | --- | --- | --- | --- |
| Comparison |  | Z | P.unadj | P.adj | Z | P.unadj | P.adj |
| M | N | -0.804078 | 0.421 | 1.000 | -1.1911933 | 0.234 | 1.000 |
| M | ORF1a | -6.5913858 | < 0.001 | **< 0.001** | -4.3091874 | < 0.001 | **< 0.001** |
| N | ORF1a | -5.7873077 | < 0.001 | **< 0.001** | -3.1179941 | 0.002 | 0.046 |
| M | ORF1b | -3.5232883 | < 0.001 | **0.012** | -4.6389208 | < 0.001 | **< 0.001** |
| N | ORF1b | -2.7192103 | 0.007 | 0.144 | -3.4477275 | 0.001 | **0.015** |
| ORF1a | ORF1b | 3.06809744 | 0.002 | 0.056 | -0.3297334 | 0.742 | 1.000 |
| M | ORF3a | -0.5484495 | 0.583 | 1.000 | -0.595282 | 0.552 | 1.000 |
| N | ORF3a | 0.25562853 | 0.798 | 1.000 | 0.59591129 | 0.551 | 1.000 |
| ORF1a | ORF3a | 6.04293627 | < 0.001 | **< 0.001** | 3.71390542 | < 0.001 | **0.006** |
| ORF1b | ORF3a | 2.97483883 | 0.003 | 0.073 | 4.04363881 | < 0.001 | **0.002** |
| M | ORF6 | -0.0066613 | 0.995 | 0.995 | -0.3089677 | 0.757 | 1.000 |
| N | ORF6 | 0.79741668 | 0.425 | 1.000 | 0.88222558 | 0.378 | 1.000 |
| ORF1a | ORF6 | 6.58472442 | < 0.001 | **< 0.001** | 4.00021971 | < 0.001 | **0.002** |
| ORF1b | ORF6 | 3.51662698 | < 0.001 | **0.012** | 4.3299531 | < 0.001 | **< 0.001** |
| ORF3a | ORF6 | 0.54178815 | 0.588 | 1.000 | 0.28631429 | 0.775 | 1.000 |
| M | ORF7a | -1.6364667 | 0.102 | 1.000 | 0.00377557 | 0.997 | 1.000 |
| N | ORF7a | -0.8323887 | 0.405 | 1.000 | 1.19496889 | 0.232 | 1.000 |
| ORF1a | ORF7a | 4.95491909 | < 0.001 | **< 0.001** | 4.31296302 | < 0.001 | **< 0.001** |
| ORF1b | ORF7a | 1.88682164 | 0.059 | 1.000 | 4.64269641 | < 0.001 | **< 0.001** |
| ORF3a | ORF7a | -1.0880172 | 0.277 | 1.000 | 0.5990576 | 0.549 | 1.000 |
| ORF6 | ORF7a | -1.6298053 | 0.103 | 1.000 | 0.31274331 | 0.754 | 1.000 |
| M | ORF8 | -0.2756125 | 0.783 | 1.000 | 0.00629262 | 0.995 | 1.000 |
| N | ORF8 | 0.52846549 | 0.597 | 1.000 | 1.19748594 | 0.231 | 1.000 |
| ORF1a | ORF8 | 6.31577323 | < 0.001 | **< 0.001** | 4.31548007 | < 0.001 | **0.001** |
| ORF1b | ORF8 | 3.24767579 | 0.001 | **0.031** | 4.64521345 | < 0.001 | **< 0.001** |
| ORF3a | ORF8 | 0.27283696 | 0.785 | 1.000 | 0.60157465 | 0.547 | 1.000 |
| ORF6 | ORF8 | -0.2689512 | 0.788 | 1.000 | 0.31526036 | 0.753 | 1.000 |
| ORF7a | ORF8 | 1.36085415 | 0.174 | 1.000 | 0.00251705 | 0.998 | 0.998 |
| M | S | -2.8735311 | 0.004 | 0.097 | -2.3660258 | 0.018 | 0.396 |
| N | S | -2.0694531 | 0.039 | 0.732 | -1.1748325 | 0.240 | 1.000 |
| ORF1a | S | 3.71785465 | < 0.001 | **< 0.001** | 1.94316163 | 0.052 | 0.988 |
| ORF1b | S | 0.6497572 | 0.516 | 1.000 | 2.27289502 | 0.023 | 0.484 |
| ORF3a | S | -2.3250816 | 0.020 | 0.401 | -1.7707438 | 0.077 | 1.000 |
| ORF6 | S | -2.8668698 | 0.004 | 0.095 | -2.0570581 | 0.040 | 0.794 |
| ORF7a | S | -1.2370644 | 0.216 | 1.000 | -2.3698014 | 0.018 | 0.409 |
| ORF8 | S | -2.5979186 | 0.009 | 0.197 | -2.3723184 | 0.018 | 0.424 |

